# Stock-production functions resembling Beverton-Holt

**DOI:** 10.1101/714956

**Authors:** Peter Fritz Baker

## Abstract

Researchers dissatisfied with the performance of the Beverton-Holt model, in contexts where “Beverton-Holt-like” behavior is expected, have introduced a plethora of alternative model forms. This paper presents a formalization of what constitutes “Beverton-Holt-like behavior” which includes many of these forms, and shows that the class of functions so defined has a coherent and non-trivial mathematical theory. Data from the stock production database assembled by Ransom Myers is used to illustrate why such generalizations have been sought in the first place, and to highlight the difficulties in choosing between model forms on purely empirical grounds. Special attention is given to a parametric family of functions within this class, here called “*θ*-BH” functions. These functions cover a broad range of shapes, including both the Beverton-Holt and hockey stick functions, and share useful properties with these two widely-used models.

## 1 Introduction

The stock-production approach to population modeling is to view abundance at some life-stage (e.g., recruitment to the ocean fisheries) as a function of abundance at some antecedent stage (e.g., the number of parent spawners). If *X* denotes the abundance of the parent stock, and *Y* the resulting production, the model is that *Y* = *F* (*X*), where the function *F* is called a “stock-production function” (or “stock-production relation”, or “stock-production law”). Although abundances are often expressed as numbers of individuals (and hence as whole numbers), it is usual to treat *X* and *Y* as continuously variable quantities, and *F* as a continuous (and even piece-wise smooth) function.

A stock-production function summarizes, and could in principle could be derived from, a deep analysis of the ecology of an organism. In practice, it is more common to use a stock production function as a *substitute* for such an analysis. That is, one argues on qualitative grounds that the “true” relationship should be at least approximately of a certain algebraic form, and selects parameters for this form by some process that side-steps the need for detailed theories of individual behavior, bioenergetics, and so on.

One commonly used form is the “Beverton-Holt” function [1]:

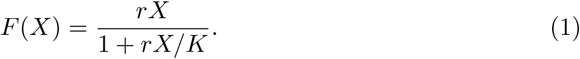

This form is appealing for a number of reasons, not least of which is that it has only two parameters, both of which can be given plausible interpretations as “real” quantities (this will be considered more carefully in Sections 4 and 5). It has become one of the most commonly used of all stock-production models.

The Beverton-Holt function has also been frequently criticized, however. Aside from reservations about the usefulness of stock-production theory in general, a recurrent complaint is that theory or empirical data suggest that Beverton-Holt has the wrong *qualitative* behavior for the situation in hand.

For example, while it is trivially true that there must be *some* theoretical upper limit on the population, this might not have any practical meaning for the population under consideration, which might be better described by the Shepherd function *F* (*X*) = *rX/*(1 + *bX*^*γ*^) with *γ* < 1 [2]. Or one might expect production to attain a maximum at some finite stock and then decrease, like the Ricker function 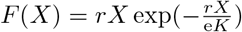 [3]. If Allee effects are a concern, the Thompson function 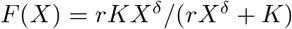 might be appropriate [4]. Some of the possibilities are illustrated in Fig 1.

**Fig 1.**
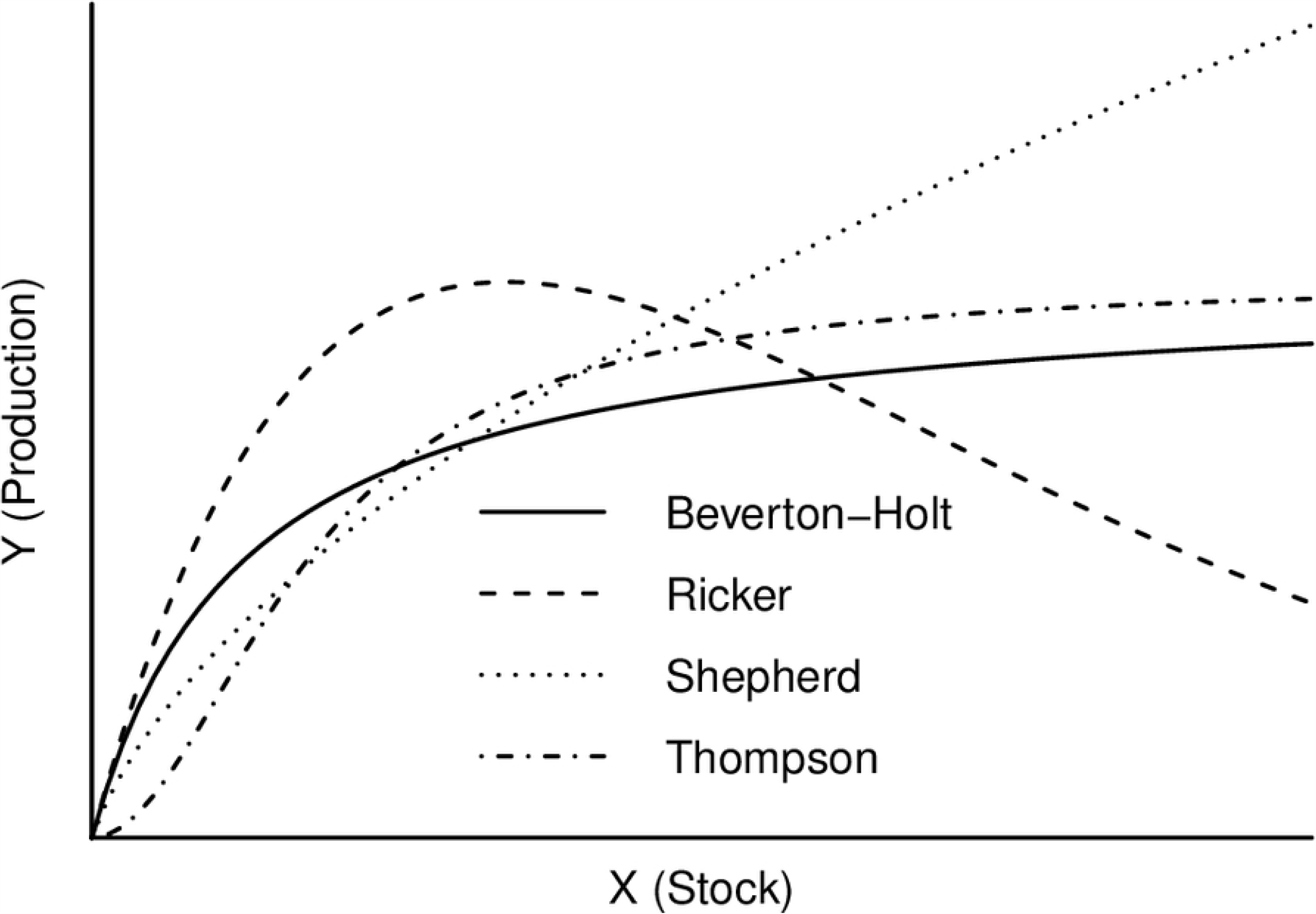
Some frequently used stock-production relationships.

This paper is concerned with a different objection: Fitting data to Beverton-Holt functions can produce results which are unsatisfactory even when the data appear “Beverton-Holt-like.” To understand this objection, and how it might be addressed, it is necessary to understand which features of the Beverton-Holt function are essential to the conceptual model underlying its adoption, and which are accidental to the particular algebraic form as Eq (1).

## 2 Beverton-Holt-like functions and r-K functions

This section formalizes what it means to say that a stock-production function “resembles” or “generalizes” the Beverton-Holt function, and discusses basic properties of the functions so defined.

### 2.1 Definitions

In many situations, biological considerations lead to the expectation that production should be nearly proportional to stock when habitat is freely available, and nearly constant when some dimension of habitat is fully utilized. In particular, this is the case when population dynamics is driven principally by contest competition [5, 6]. These qualitative expectations can be formalized as constraints on the mathematical form of a stock-production function *Y* = *F* (*X*):

I. For some 0 *< r < ∞*, the graph of *F* approaches the line *Y* = *rX* as *X* goes to 0.
II. For some 0 *< K < ∞*, the graph of *F* approaches the line *Y* = *K* as *X* goes to *∞*. Conventionally, *r* is interpreted as an “intrinsic rate of increase” or a “density-independent survival,” and *K* as the ultimate “carrying capacity” of the habitat (this will be considered more carefully in Section 4). These are typically are the only properties that are justifiable *a priori*, so *F* should be otherwise unremarkable. A simple formulation of this is:
III. *F* (*X*) is non-decreasing and concave.

A more stringent notion of “unremarkability” is explored in S1 Appendix.

For the rest of this paper, continuous functions satisfying the conditions (I), (II), and (III) will be called *Beverton-Holt-like*. It is mathematically convenient to exclude the case when stock (and hence production) is exactly zero, so Beverton-Holt-like functions will be continuous functions on the positive real numbers.

Continuous functions on the positive real numbers satisfying the conditions (I) and (II) will be called *r-K functions*.

### 2.2 Standard form and homogeneous families

The limiting values associated with an r-K function *F* will be denoted **r**(*F*) and **K**(*F*). The (signed) area of the “wedge” between the graph of *F* and the line *y* = **K**(*F*) will be called the *defect* of *F*, and written defect(*F*); this is one way of measuring how quickly the function approaches its limit. Fig 2 shows a generic Beverton-Holt-like function.

**Fig 2.**
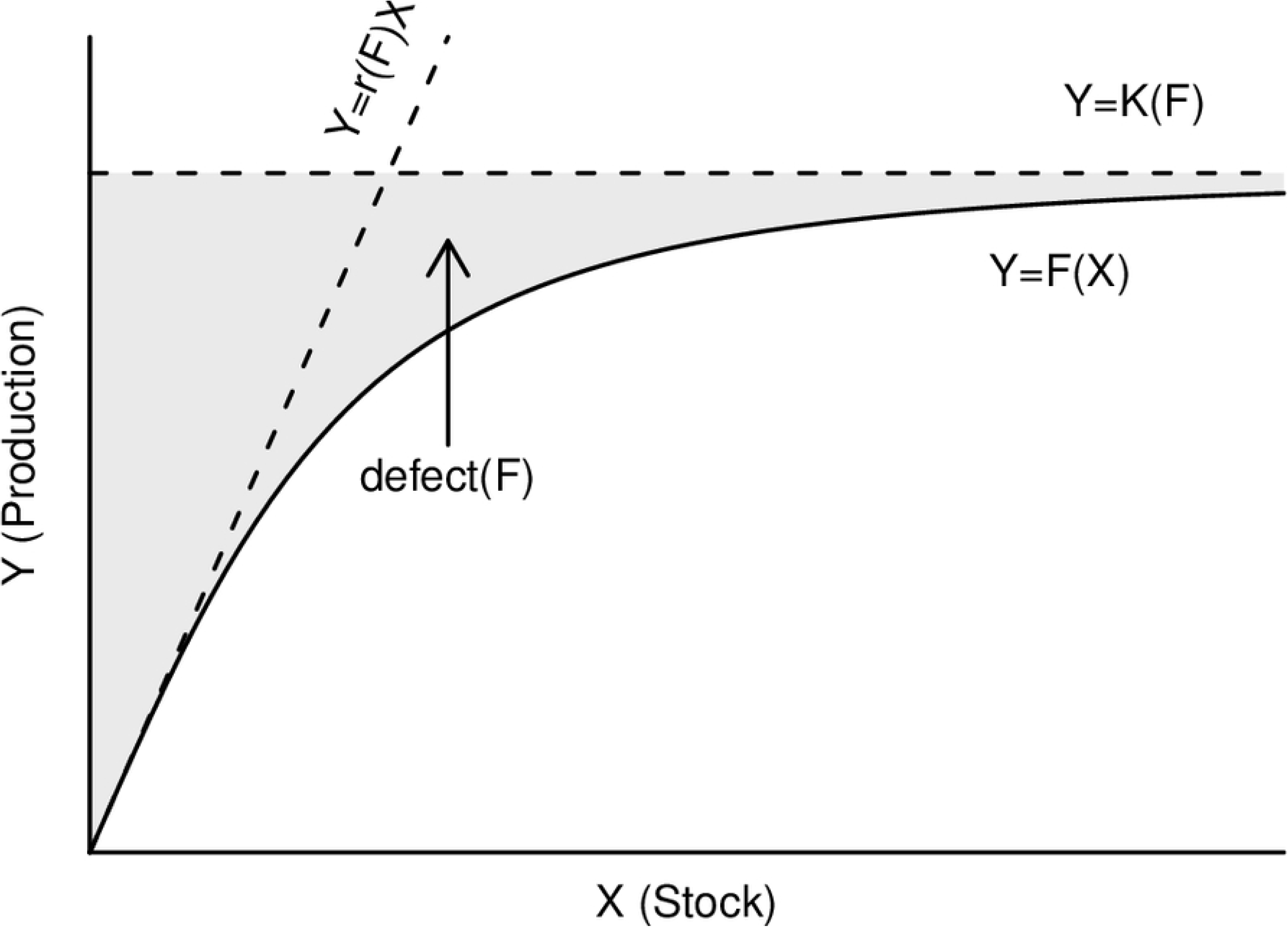
A generic Beverton-Holt-like function (solid line), with its defining asymptotes (dashed lines) and defect (shaded area).

An r-K function *f* will be said to be in *standard form* if **r**(*f*) = **K**(*f*) = 1. If *f* is in standard form, then the function *F* (*X*) = *Kf* (*rX/K*) is an r-K function with **r**(*F*) = *r* and **K**(*F*) = *K*. Conversely, every r-K function *F* is associated in this way with a unique function in standard form, which will be called the *standard form of F*.

A family 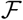 of r-K functions will be called *homogeneous* if all its members have the same standard form. This common form will be called the *standard form of* 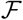.

A family 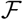 of r-K functions will be called *full* if every possible combination of *r* and *K* is represented by at least one member of 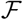.

In practice, Beverton-Holt-like functions encountered in the stock-production literature seem to fall naturally into homogeneous families. This is partially explained by the following observation:

**Proposition 1.**

*Suppose 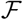 is a full family of r-K functions whose members are uniquely characterized by their r and K values. Suppose also that 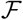 is closed with respect to rescaling; that is, for any F in 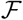 and any positive numbers a and b, the function aF* (*bX*) *is also in 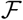. Then 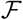 is homogeneous*.

*Proof.* Let *f* be the unique member of 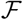 with **r**(*f*) = 1, **K**(*f*) = 1. If *F* is any other function in 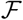, say with **r**(*F*) = *r*, **K**(*F*) = *K*, then *F* (*X*) and *Kf* (*rX/K*) are two members of 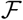 with the same *r* and *K* values; by assumption, these must coincide.

### 2.3 Examples of Beverton-Holt-like functions

Some examples of Beverton-Holt-like functions are collected in Table 1 in standard form, together with their defects. Fig 3 shows a sampling of these.

**Table 1.** Examples of Beverton-Holt-like functions in standard form. To express these in the original units for stock and production, apply the substitutions *x* = *rX/K*, *y* = *Y / K*, and multiply the defects by *K*^2^*/r*.

| Name | Reference | Standard form* | Defect<sup>†‡</sup> |
| --- | --- | --- | --- |
| hockey stick | [7] | $y = \min(x, 1)$ | $\frac{1}{2}$ |
| Beverton-Holt | [1] | $y = \frac{x}{1+x}$ | ∞ |
| Skellam | [8] | $y = 1 - e^{-x}$ | 1 |
| θ-BH | [9] | $y = \frac{x}{(1+x^\theta)^{1/\theta}}$ | $\begin{cases} \infty & \text{if } 0 < \theta \leq 1 \\ \frac{1}{\theta} B(\frac{2}{\theta}, 1 - \frac{1}{\theta}) & \text{if } 1 < \theta < \infty \end{cases}$ |
| negative power | [6] | $y = 1 - (1 + \frac{x}{\lambda})^{-\lambda}$ | $\begin{cases} \infty & \text{if } 0 < \lambda \leq 1 \\ \frac{\lambda}{\lambda-1} & \text{if } 1 < \lambda < \infty \end{cases}$ |
| positive power | (see text) | $y = 1 - (1 - \frac{x}{\lambda})_+^\lambda$ | $\frac{\lambda}{\lambda+1}$ for $1 \leq \lambda < \infty$ |
| logistic hockey stick | [7] | $y = 1 - \log_\lambda(1 + (\lambda - 1)\lambda^{-\frac{\lambda}{\lambda-1}x})$ | $-\frac{\lambda-1}{\lambda(\log \lambda)^2} \text{Li}_2(\lambda)$ for $1 < \lambda < \infty$ |
| bent hyperbola | [10] | $y = \frac{2x}{x+1+\sqrt{x^2-2\lambda x+1}}$ | $\begin{cases} \infty & \text{for } -1 \leq \lambda < 1 \\ \frac{1}{2} & \text{for } \lambda = 1 \end{cases}$ |

\* $(x)_+$ is $\max(0, x)$ .
<sup>†</sup> $B(\cdot, \cdot)$ is the beta function, $B(u, v) = \int_0^1 t^{u-1}(1-t)^{v-1} dt$ [11, §6.2].
<sup>‡</sup> $\text{Li}_2(\cdot)$ is the dilogarithm function, $\text{Li}_2(u) = -\int_1^u \log(t)/(t-1) dt$ [11, §27.7].

**Fig 3.**
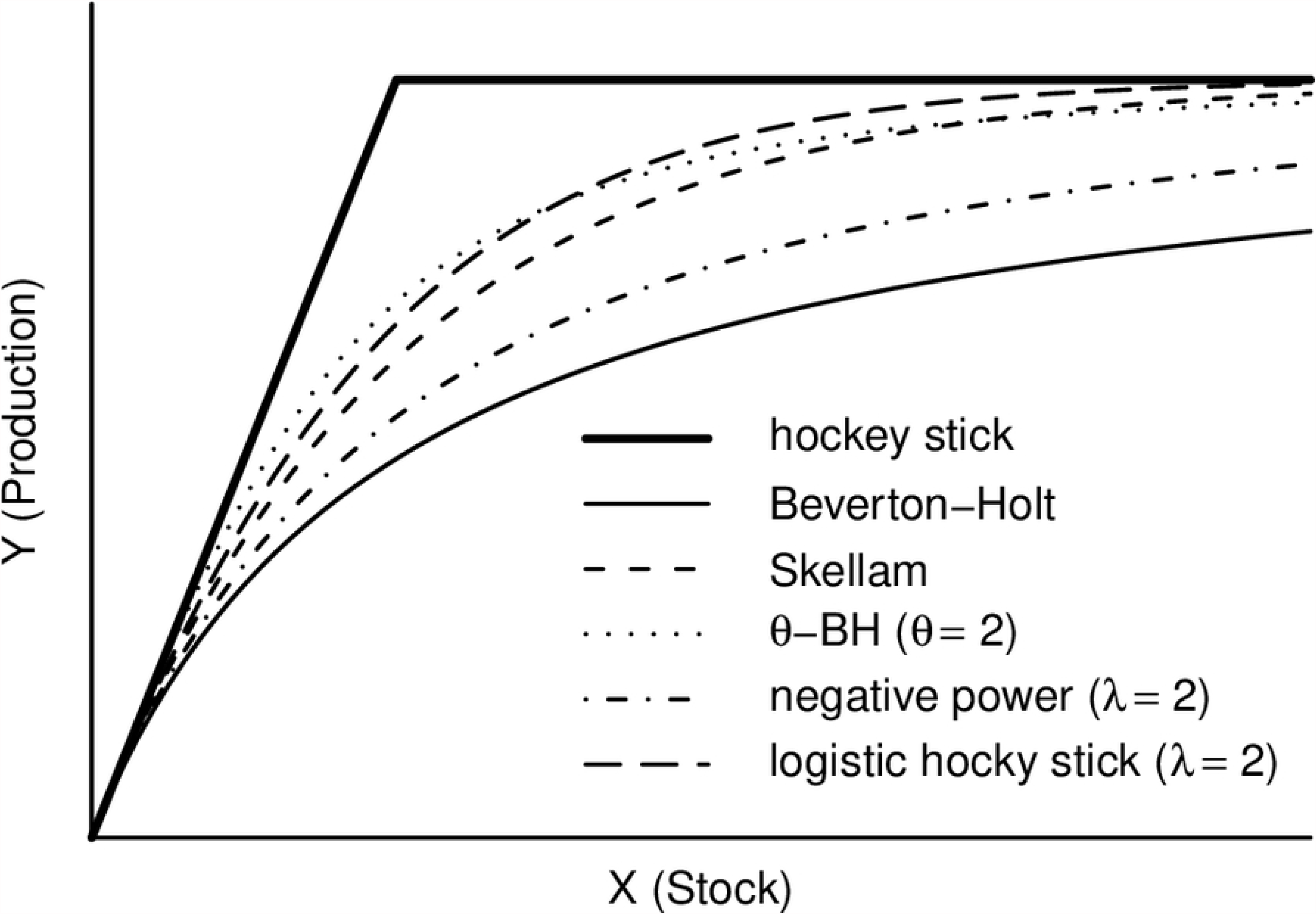
Some Beverton-Holt-like functions having the same *r* and *K*. See Table 1 for sources and algebraic forms.

Since all Beverton-Holt-like functions look more or less alike to the human eye, it is reasonable to wonder if there is any point to this proliferation of algebraic forms. This will be addressed in Sections 4 and 6; for now, it will just be noted that many of the functions in Table 1 were introduced specifically to address *practical* problems encountered with Beverton-Holt.

The hockey stick, Beverton-Holt, and Skellam functions have all been used by multiple researchers. Moreover, less familiar functions seem invariably to include one or more of these as a special or limiting cases. It is obvious that any Beverton-Holt-like function is bounded above by the hockey stick of the same *r* and *K*, and it is shown in and it is shown in S1 Appendix that any Beverton-Holt-like function which is “sufficiently boring” (in a technical sense explained in the Appendix) is bounded above by the Skellam of the same *r* and *K*.

The name “*θ*-BH” was coined for the present paper, in analogy to the *θ*-logistic differential equation with which it is closely related (Section 3.3). [9] calls it the “*δ*-power B&H”, and derives it from a size-based recruitment model. It makes a fleeting appearance in [12] without acquiring a name there. Despite its simplicity, it has not seen much use as a stock-production model. The *θ*-BH functions can be obtained from Beverton-Holt by a simple change of variables: *Y* is a *θ*-BH function of *X* if and only if *Y*^*θ*^ is a Beverton-Holt function of *X*^*θ*^.

The negative power distribution is derived in [6] as a mixture of Skellam functions. Mixtures of Beverton-Holt-like functions will be discussed more generally in Section 2.5. The positive power distribution is a trivial variant of this, but I haven’t been able to find a case in which it has been used as a stock-production relation (perhaps because it is not easily fitted by the usual least-squares method).

The logistic hockey stick is presented in [7] as a function in three parameters *α*, *µ*, and *θ*, most naturally expressed as a cumulative integral:

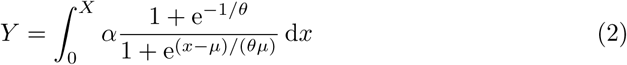

The standard form of this turns out to depend only on *θ*, so this is a natural shape parameter; when working with the standard form, it is convenient to use *λ* = e^1/*θ*^ + 1, resulting in the (admittedly bizarre) expression in Table 1. The natural domain of *λ* from this derivation is 1 *< λ < ∞*, with the limiting cases *λ →* 1 and *λ → ∞* yielding the Skellam and hockey stick, respectively. The algebraic expression in the table continues to describe a Beverton-Holt-like function for the larger interval 0 < *λ* < ∞, with a removeable singularity at *λ* = 1.

The bent hyperbola is the specialization of the more general bent hyperbola of [13] to the context of Beverton-Holt-like functions. It is presented in [10] as a function in three parameters *β*, *S^∗^*, and *γ*:

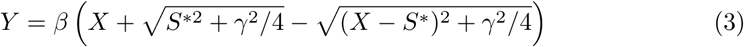

The standard forms of these turn out to depend only on the quantity 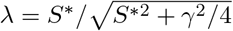, which is taken as the shape parameter here. The natural domain of *λ* from this derivation is the closed unit interval *−*1 *≤ λ ≤* 1, with the limiting cases *λ* = *−*1 and *λ* = 1 yielding the Beverton-Holt and hockey stick, respectively. The algebraic expression in the table continues to describe a Beverton-Holt-like function for the larger interval −∞ < *λ* ≤ 1.

The bent hyperbola reveals a limitation of the defect as a measure of distance from the hockey stick: although *λ* = 1 is the hockey stick, and the convergence as *λ →* 1 is uniform, the defect is infinite for all *λ <* 1.

### 2.4 Cumulative distributions

The definition of Beverton-Holt-like functions given in 2.1 does not include any explicit smoothness requirements. However, it is a standard mathematical result that any concave function must be quite well behaved—for example, locally Lipschitz continuous and differentiable at all but countably many points [14].

In particular, any Beverton-Holt-like *F* can be written as a cumulative integral:

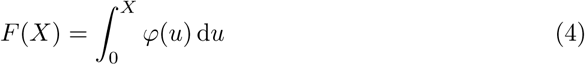

where *φ* is a monotone non-increasing function on (0, ∞) with lim_*u→*0_ *φ*(*u*) = **r**(*F*) and 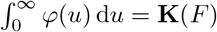. Conversely, any non-negative, monotone non-increasing function which is bounded above and integrable on (0, ∞) gives rise to a Beverton-Holt-like function via Eq (4).

If *F* is in standard form, *φ* is a probability density on (0, ∞). This gives an interesting interpretation of the defect: if *φ* is the density function associated with an *f* in standard form, then

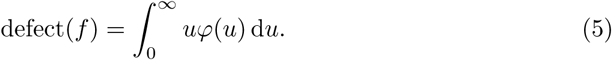

That is, defect(*f*) is the expected value for the probability distribution associated with *f*.

The Skellam function is associated in this way with the standard exponential distribution [15, Chapter 19], and the hockey stick with the uniform distribution on (0, 1) [16, Chapter 26].

Other functions from Table 1 correspond to less familiar distributions: the positive power function is associated with a (non-standardized) beta distribution with *p* = 1*, q* = *λ* [16, Chapter 25]; the *θ*-BH and negative power functions are associated with the special cases *p* = *θ, q* = 1 and *p* = 1*, q* = *λ*, respectively, of the generalized *t*-distribution of [17].

The density corresponding to the logistic hockey stick for *λ >* 1 does not seem to have found much use by statisticians to date. For 0 *< λ <* 1, however, this is the Exponential-Logarithmic distribution of [18] with 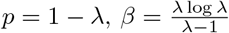.

### 2.5 Mixtures

If *F* and *G* are Beverton-Holt-like functions and *a, b* are non-negative constants, *aF* + *bG* is also Beverton-Holt-like.

More generally, let (Ω*, µ*) be any measure space, and let {*F*_*ω*_}_*ω*∈Ω_ be a family of Beverton-Holt-like functions such that the map 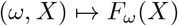 is

Ω *×* (0, ∞)-measurable. If both 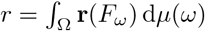 and 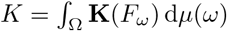 are finite, the mixture

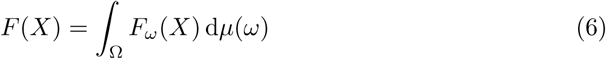

is Beverton-Holt-like, with **r**(*F*) = *r* and **K**(*F*) = *K*.

Considerations of habitat heterogeneity lead naturally to such mixtures. This will be pursued a bit further in S1 Appendix.

The “negative power” function, which includes both Skellam and Beverton-Holt as limiting cases, can be exhibited as a continuous mixture of Skellam functions:

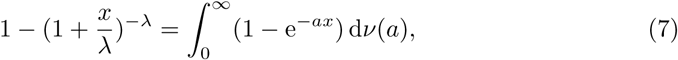

where *ν* is a gamma distribution with expected value 1:

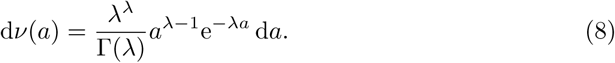

If the term 1 + *x/λ* is modified to (1 + *x/λ*)_+_, the negative power function continues to be an r-K function for *negative* values of the parameter *λ*, and is Beverton-Holt-like when *λ ≤ −*1. This “positive power” function has some charm: it has the interesting (and biologically natural?) property of actually attaining the production capacity at a finite stock. It is not, however, a mixture of Skellam functions. (This will follow from results in S1 Appendix.)

### 2.6 Composition

If *Y* is a Beverton-Holt-like function of *X*, and *Z* is a Beverton-Holt-like function of *Y*, then *Z* is also a Beverton-Holt-like function of *X*. That is, compositions of Beverton-Holt-like functions are again Beverton-Holt-like. In particular, models obtained by iterating Beverton-Holt-like functions cannot produce “interesting” population dynamics, as iteration of Ricker functions famously can [19].

Compositions of true Beverton-Holt functions are also true Beverton-Holt functions, whose parameters are simple combinations of the parameters of the constituents. Specifically, if *F* and *G* are Beverton-Holt, their composition *G* ◦ *F* is the Beverton-Holt with parameters

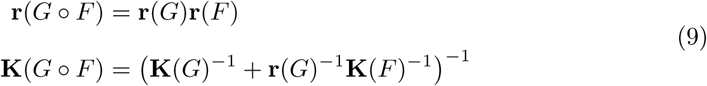

This property is sometimes convenient for simulation modeling [20]. It is exploited systematically, for example, in the EDT framework of [21].

The Beverton-Holt family is not the only family with a “composition law”, however. If *F* and *G* are *θ*-BH for the same value of *θ*, *G* ◦ *F* is also *θ*-BH for this *θ*, with

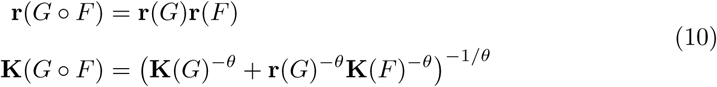

Moreover, if *F* and *G* are hockey sticks, *G* ◦ *F* is also a hockey stick, with

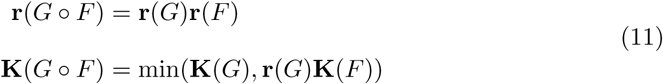

Since the limit as *θ* → ∞ of the *θ*-BH functions having given *r* and *K* parameters is the hockey stick with these parameters, hockey sticks can be thought of as ∞-BH functions; with this convention, Eqs (9) and (11) are both special cases of Eq (10).

It turns out that these are the *only* examples of full homogeneous families with composition laws:

#### Theorem 2.

*Suppose that 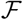 is a full homogeneous family of r-K functions, and suppose that 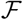 is closed under composition. Then there is some 0 < θ ≤ ∞ such that 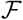 is the family of θ-BH functions*.

Since this is a mathematical fact, rather than a biological one, the demonstration is relegated to S2 Appendix.

## 3 Stock-production and continuous dynamical systems

Stock-production functions can arise as discrete dynamical systems derived from continuous dynamical systems. In particular, the Beverton-Holt function has a natural association with the widely used logistic differential equation for the evolution of populations over time. Furthermore, this association persists even when the parameters of the equation are allowed to vary with time.

This section will describe this connection, and show that the *θ*-BH functions are associated in the same way with *θ*-logistic differential equations.

### 3.1 Differential equations and stock-production

An approach to population modeling with a very long history is to consider abundance as a continuous quantity, varying continuously in time, and satisfying a first-order differential equation derived from consideration of changes in arbitrarily short time increments:

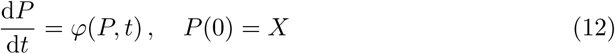

for the abundance *P*, where *X* is the initial population and *φ* is a continuous function from (0, ∞) *×* [0, ∞) into (*−∞, ∞*).

It is shown in standard textbooks on differential equations that, under mild assumptions on *φ*, Eq (12) has a unique solution for any *X >* 0, at least on some non-trivial interval containing *t* = 0 [22]. If there is some interval [0*, T*_max_) which works for *all X >* 0, then any *T* strictly between 0 and *T*_max_ gives rise to a stock-production relation *F* between *X* = *P* (0) and *Y* = *P* (*T*).

Stock-production functions derived in this way are necessarily monotone, as noted in [23, §3.1.2]. S3 Appendix explores conditions under which they are Beverton-Holt-like.

### 3.2 The exponential and logistic equations

Perhaps the oldest population model of all is the differential equation

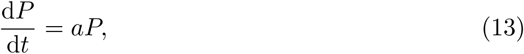

with *a* constant. This appears (implicitly) already in the pioneering work of Graunt on human demographics [24]. The solution, *P* (*t*) = *P* (0)e^*at*^, gives rise to the stock-production function *Y* = *rX*, where *r* = e^*aT*^.

Malthus observed that exponential growth cannot persist indefinitely, and must therefore be modified by some kind of density dependence [25]. An early mathematical formulation of this is the logistic differential equation of Verhulst [26],

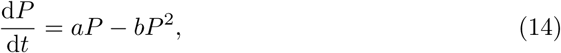

with *a* and *b* constant, *b >* 0. The solution to this is the classic logistic function

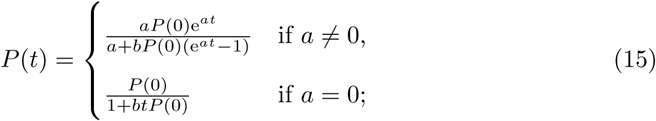

the corresponding stock-production function is therefore the Beverton-Holt function with parameters

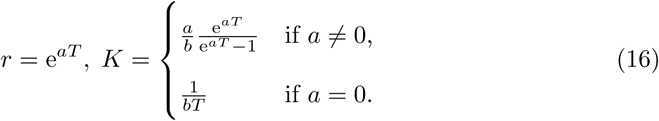

### 3.3 The *θ*-logistic equation

Typical derivations of Eq (14) have an *ad hoc* flavor, starting from the desired qualitative behavior of *φ* and simply taking the “simplest” form that works [26, 27].

Just what “simplicity” means in this context is not entirely clear, however, and alternatives have been considered from a very early date. In particular, the models

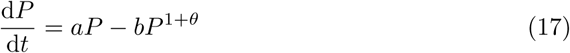

for *θ >* 0 already appear in Verhulst’s 1838 paper [26, page 116].

Eq (17) is sometimes called the “*θ*-logistic” equation. In population-modeling contexts, it has been called the “Richards equation” after its appearance in [28].

The substitution *Q* = *P*^*θ*^ reduces (17) to the ordinary logistic equation d*Q*/d*t* = *aθQ − bθQ*^2^, so the associated stock-production model is the *θ*-BH function with parameters

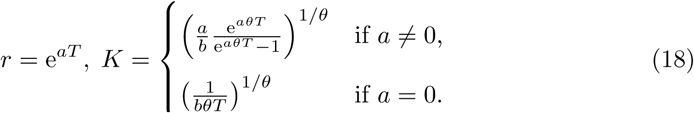

### 3.4 Time-varying parameters

The models (13), (14), and (17) are “autonomous,” that is, the function *φ* does not depend explicitly on *t*. Since environmental conditions can be expected to vary over time, it is natural to consider generalizations of these models in which the coefficients are functions of *t*. From the stock-production point of view, this turns out to add nothing new.

Specifically, the general solution to (17), considered as a possibly *non*-autonomous equation, can be written as

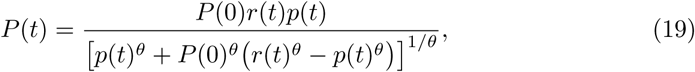

where *p*(*t*) is the particular solution with *p*(0) = 1 and 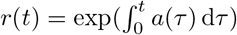. If *b*(*t*) ≡ 0 on the interval [0*, T*], the corresponding stock-production function is simply *Y* = *r*(*T*)*X*. Otherwise it is *θ*-BH, with parameters

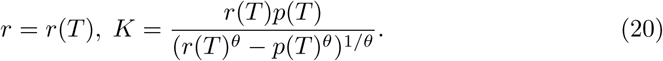

### 3.5 Another characterization of *θ*-BH functions

The stock-production functions associated with a given *φ* as in Section 3.1 for different choices of *T*, form a one-parameter family.

More generally, let *φ* be a continuous function from (0, ∞) *×* [0, ∞) into (*−∞, ∞*), and suppose that there is some 0 *< T*_max_ *≤ ∞* such that, for each 0 *≤ s < T*_max_ and each 0 *< X < ∞*, the initial value problem

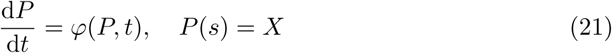

has the unique solution *F*^*s,t*^(*X*), *s ≤ t < T*_max_. Then 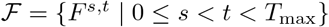 is a two-parameter family of stock-production functions, degenerating to a one-parameter family in the autonomous case (since then *F*^*s,t*^ = *F* ^0*,t−s*^).

In the cases considered in Sections 3.2–3.4, this family was homogeneous. This turns out to be a very special property:

#### Theorem 3.

*Let 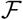 be as above. If the F ^s,t^ are all r-K functions having the same standard form, this form is the standard θ-BH for some* 0 *< θ < ∞*.

This result is closely related to Theorem 2. A proof is given in S2 Appendix.

## 4 Beverton-Holt-like functions as population models

It is hard to avoid the suspicion that all this mathematics is out of proportion to the original problem—that the proliferation of algebraic forms for the stock-production function is more a matter of scholastic hair-splitting than practical biology.

Such doubts are only exacerbated by the fact that such functions are typically used in a crudely empirical way. The usual goal is simply to *describe* a data set, with no pretense that the fitted form is anything other than a conveniently simple approximation to an inconveniently complex reality.

This section and the next will try to explain why so many “alternatives to” or “generalizations of” the basic Beverton-Holt law have nonetheless been proposed over the years.

### 4.1 Physical interpretation of model parameters

In the conceptual model of Section 2, the *r* and *K* parameters correspond to “real” quantities:

- *r* is the “intrinsic rate of increase” one would expect to see in the absence of crowding—a measure of *habitat quality*.
- *K* is the “carrying capacity” or maximum production potential—a measure of *habitat quantity*.

There are thus *two* sets of “*r*” and “*K*” parameters present when a Beverton-Holt-like model is considered: values describing the habitat, and values describing the population dynamics.

Some applications of stock-production modeling rely on identifying the two. This can be done in either direction:

- One can attempt to obtain information about physical habitat from population data. For example, parameters obtained by fitting a stock-production model to population data may be used as estimates of physical values, in the course of setting harvest levels [29] or estimating extinction risks [30].
- One can attempt to obtain information about population dynamics from habitat data. For example, stock-production models parameterized with physical values (from survival experiments, habitat mapping, etc.) may be used to game management or restoration alternatives [20].

What makes this identification dangerous is that the *r* and *K* parameters *by definition* concern properties of the fitted curve at the fringes of the data (strictly speaking, outside any *possible* range of data). Estimating intrinsic productivity or ultimate carrying capacity parameters from passively-observed stock-production data, by fitting *any* kind of stock-production model, is always extrapolation; such values may be driven as much by the algebraic form of the model as by the data.

### 4.2 Beverton-Holt as a Beverton-Holt-like function

A series of papers by Ransom A. Myers, Nicholas J. Barrowman, and others [7, 31, 32] have developed an empirical case that use of the Beverton-Holt model to infer physical-habitat parameters from population dynamics tends to overestimate both *r* and *K*, which can lead managers to overestimate the robustness of populations with respect to exploitation or habitat loss.

Of course, it is to be expected that using the wrong model will produce incorrect results—the Beverton-Holt function is not special in this regard. However, if it is *typical* for “true” stock-production curves to sit above the Beverton-Holt curve having the same asymptotes, then fitting Beverton-Holt functions will *typically* over-estimate *r* and *K*. And this is precisely what is asserted in in the cited papers.

At the root of these charges is the very slow rate at which the Beverton-Holt function rises toward its asymptote, which corresponds to a seemingly inefficient use of resources. For example, Eq (1) predicts that even when the habitat is 100% over-seeded (in the sense that *X* = 2*K/r*, the stock which would yield a production of 2*K* in the absence of density-dependence), fully one-third of the productive potential of the habitat will remain unexploited (in the sense that *Y* = 2*K/*3). Whether this is reasonable or not of course depends on how organisms actually interact with the habitat, and with one another, in the situation at hand. But as a generic assumption, to be used in the absence of population-specific detail, it is at least open to challenge.

One measure of just how slowly the Beverton-Holt function approaches the horizontal asymptote is that the “wedge” between the curve and the asymptote has infinite area. In the notation of Table 1, defect(*F*) = *∞* when *F* is a Beverton-Holt function. Since real populations are always finite, it is not clear how to interpret this. However, it is noteworthy that most of the Beverton-Holt-like functions that have been proposed over the years as alternatives to Beverton-Holt have finite defect. If a Beverton-Holt-like function in standard form is interpreted as the cumulative function of a probability density, as in Section 2.4, the defect is finite whenever this density has an expected value; Section 6 of Web Appendix A gives another heuristic under which functions of infinite defect might be considered “unreasonable.”

## 5 Beverton-Holt-like functions in practice

This section will make the discussion of Section 4 more concrete by considering some actual data. Because the papers of Myers et al. are so cogent, and because these authors have made their data publicly available with the express desire to support “meta-analytic methods to combine results across many populations” [33], they seem particularly well suited to the present purpose. The focus will be on populations of Coho salmon *Oncorhynchus kisutch* discussed in [7].

### 5.1 Coho salmon data

Fig 4 shows all the sets of stock-production data from the Myers database [34] for Coho salmon, *Oncorhynchus kisutch*, in which “stock” is female spawners and “production” is outmigrant smolts, arranged very subjectively by shape.

**Fig 4.**
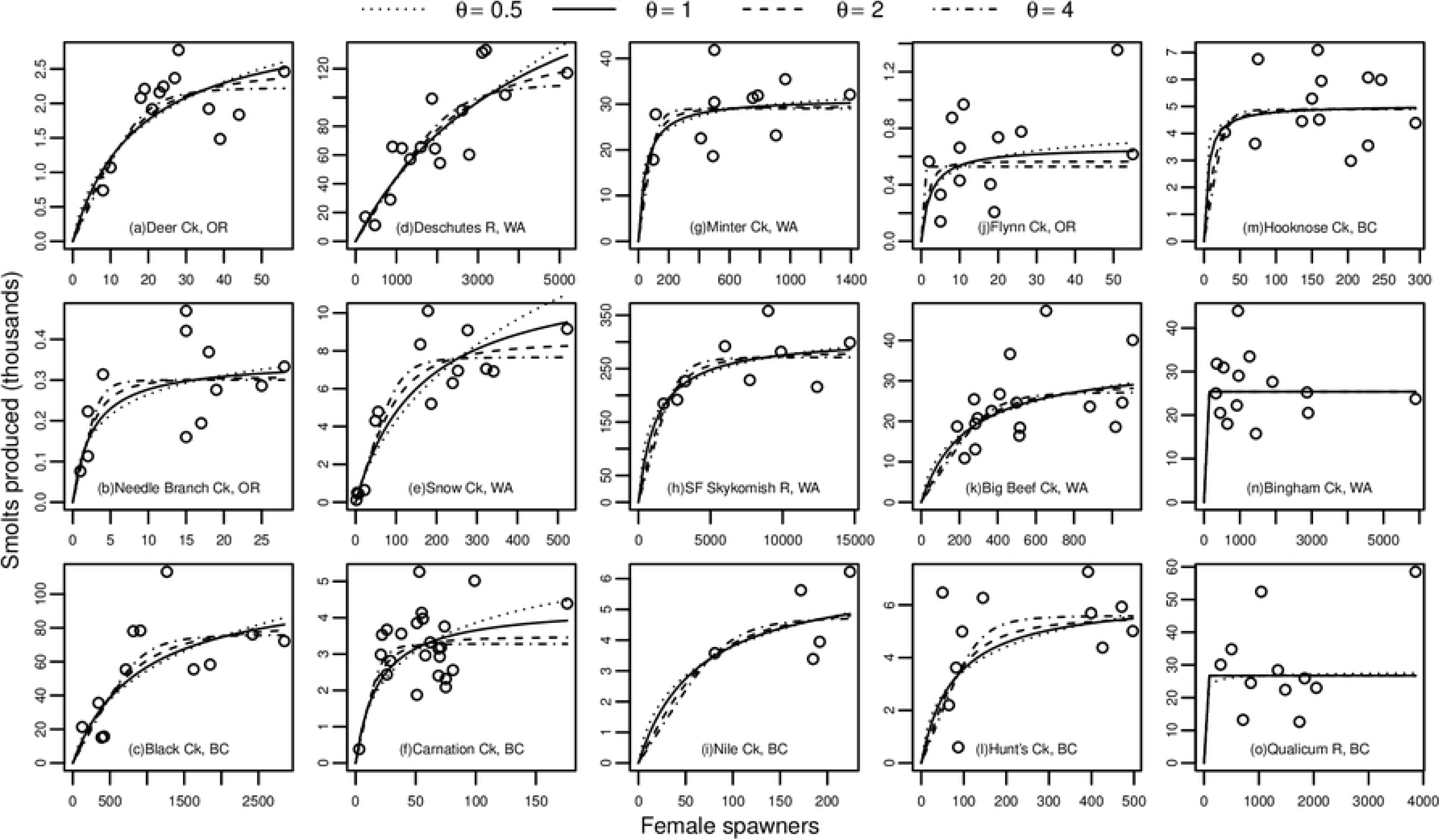
Coho salmon (*Oncorhynchus kisutch*) female-to-smolt data from the Myers database, with fitted *θ*-BH curves.

The fitted curves will be discussed below. Disregarding these for the moment, several features of these data are worth remarking.

First, all the individual datasets are quite small. The largest consists of 26 points. This is typical of stock-production data: monitoring programs that follow a consistent methodology for a quarter of a century are unfortunately (albeit understandably) rare.

Next, the data are quite noisy. Again, this is typical of stock-production data: environmental drivers (flow, water temperature, food availability) can vary dramatically from year to year, and the estimates of both stock and production usually come from sampling, with large uncertainties.

Because the datasets are small and noisy (or as Ransom Myers put it, “nasty, brutish, and short” [33]), it is unrealistic to expect the data to point unambiguously to a particular algebraic form. One might decide that a dataset seems “Beverton-Holt-ish”, or “Ricker-ish”, but it seems hopeless to choose between, say, the logistic hockey stick and bent hyperbola, on empirical grounds alone. It will be seen that there are good reasons for considering models with a degree of freedom beyond *r* and *K*, but one more “shape” parameter is probably all that can be accommodated.

Many of these particular datasets are plausibly Beverton-Holt-like. Considered in isolation, some of these sets (e.g. a–c) plausibly convey information about both *r* and *K*, without the need to assume much except a general “Beverton-Holt-like-ness.” Others suggest constraints on either *r* (d–f) or *K* (g–i), but not necessarily both. Still others (m–o) hint at possible non-Beverton-Holt-like *decreases* in production at high density.

### 5.2 Model fitting

The observed stock-production pairs (*X*_*i*_, *Y*_*i*_), 1 ≤ *i* ≤ *n*, are assumed to be related via a stock-production function *F* from some parametric family *F* (*⋅*, **p**). Since the purpose here is simply illustrative, a simple multiplicative error structure is assumed:

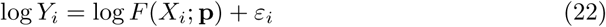

where the *ε*_*i*_ are i.i.d. normal deviates with mean 0 and variance *σ*^2^.

(Stock-production datasets are typically time-series, with *X*_*i*_ in turn a function of earlier *Y*_*j*_, *j < i*, and autocorrelation should be taken into account. Furthermore, both *X*_*i*_ and *Y*_*i*_ are usually themselves estimated from sampling data, and hence have error structures of their own. A more general framework here is state-space modeling [35].)

Eq 22 gives rise to the log-likelihood

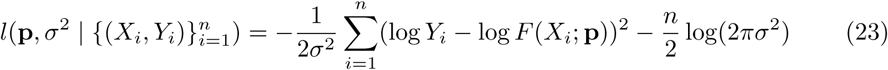

The maximum-likelihood estimate for the parameters is 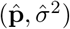, where 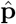 minimizes the residual sum of squares

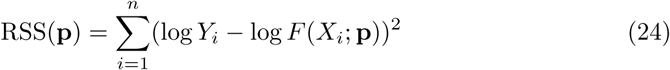

and 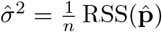.

The Bayesian Information Criterion is then

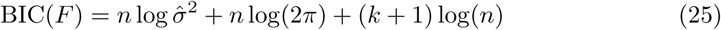

where *k* is the length of **p**. Given an alternate model form

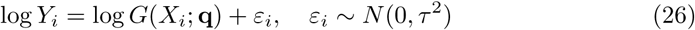

the performances of the maximum-likelihood-fitted models will be compared as

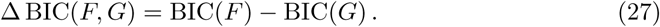

The quantity 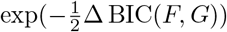 is interpretable as an evidence ratio.

The models considered will all be *θ*-BH:

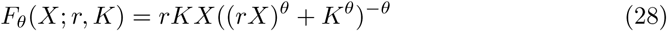

The present concern is how the performance of this model depends on *θ*. That is, rather than treating *θ* as a fitting parameter, a range of values for *θ* are considered, and the log-likelihood maximized with respect to the remaining parameters. The intention is to explore how choice of algebraic form affects the estimation of the “natural” quantities *r* and *K*, using the family of *θ*-BH functions for different values of *θ* as a convenient surrogate for the full range of possible r-K functions.

### 5.3 Results

The methods of Section 5.2 were used to fit *θ*-BH models to the Myers Coho data for a range of *θ*. Four of these fitted curves are shown in Fig 4 for each of the data sets. Fig 5 shows how the fitted parameters and BIC depend on *θ*. The ∆ BIC values are all calculated with respect to the model with *θ* = 1.

**Fig 5.**
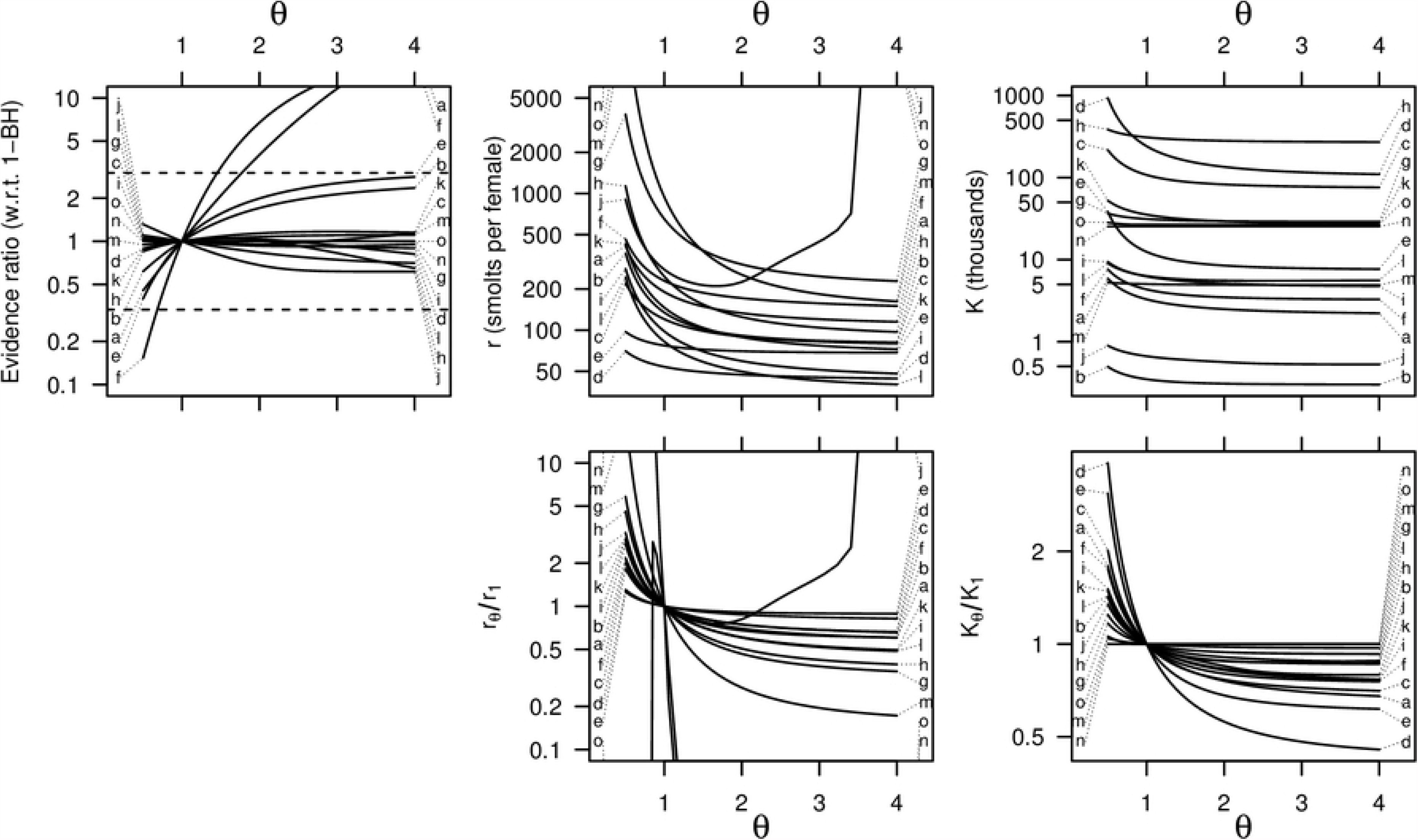
Dependence of fitted models on the shape parameter *θ*. By at least one widely-used criterion, a broad range of *θ* values are more-or-less equally compatible with the data. The fitted *r* and *K* parameters however, vary considerably between models (note the logarithmic scale).

In most cases, the fitted curves look very similar to one another. This subjective impression is supported by the evidence-ratio panel of Fig 5: for most of the datasets, the evidence ratio for the *θ*-BH model is within a factor of three of the Beverton-Holt model over a wide range of values for *θ*, and thus by a widely-used rule-of-thumb all these models are comparably well supported. The fitted values of *r* and *K*, on the other hand, depend quite strongly on *θ*.

To put it more dramatically, it is not possible to estimate *r* and *K* very well without committing to a particular model form, and the data themselves are of little help in selecting this form. Recall that this is simply a range of models consistent with the Beverton-Holt-like heuristic: as noted in Section 1, these by no means cover the universe of forms considered seriously in the stock-production literature.

Essentially the same phenomenon is analyzed in [36], where a discretized version of the *θ*-logistic differential equation discussed in Section 3 is fitted to real and simulated population time-series. These authors show that treating *θ* as a fitting parameter on par with the (appropriate analogs of) *r* and *K* is very unstable, in the sense that small perturbations of the data can result in large changes to the parameter estimates, and in the case of simulated series, that model fitting cannot reliably recover the “true” values used to generate the series in the first place.

### 5.4 Other taxa

The same patterns are seen for other taxa.

The great majority of stocks in the Myers database are assigned there to one of six orders of bony fishes: Clupeiformes, Gadiformes, Perciformes, Pleuronectiformes, Salmoniformes, and Scorpaeniformes. Of these stocks, 543 have at least five years of stock and production values. Applying the fitting procedure of Section 5.2 to these data yields the evidence-ratio curves shown in Fig 6.

**Fig 6.**
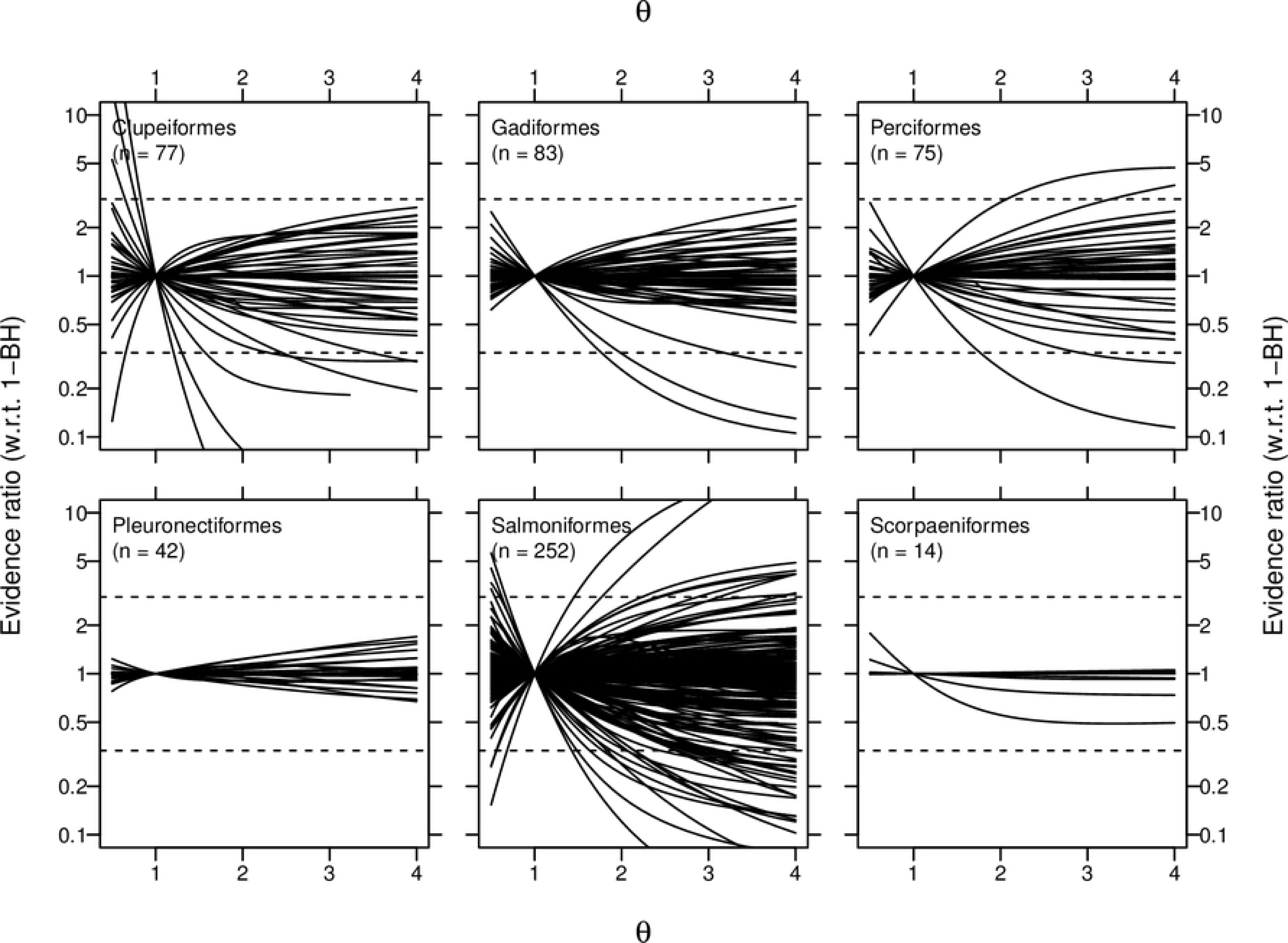
Evidence-ratio curves for *θ*-BH fits to stock-recruitment time-series for all stocks from selected orders from the Myers database.

All but a handful of these curves are monotone on the entire interval 0.5 *≤ θ ≤* 4. That is, if *θ* is treated as a parameter on a par with *r* and *K*, to be estimated by maximum likelihood, the fitting routine wants to drive the model to one of the two degenerate forms *θ → ∞* or *θ →* 0.

The curves which do have a local maximum are very flat: in only eleven cases is the evidence ratio at some intermediate *θ* more than 20% higher than at both endpoints. For 170 (31%) of the stocks, the curves are “extremely flat,” in the sense that they vary no more than 1% across the full range of *θ*.

### 5.5 Remarks

It should be emphasized that the analyses of this section are meant to illustrate some general points about stock-production relationships. A serious analysis of the Coho data from the point of view of a regional manager, say, would be much more involved. Such an analysis would require a more careful consideration of error structures, and would probably treat some or all the populations together (as recommended, for example, in [37]).

First and foremost, however, such an analysis would consider environmental covariates. There is a mistaken perception that stock-production ideas are somehow in conflict with environmental explanations for population levels (a legacy, perhaps, of otherwise forgotten ideological/philosophical debates from the early twentieth century [38]). In principle, the relationship between stock and production for any particular cohort is *determined* by the environmental circumstances of that cohort, and should vary, perhaps dramatically, from cohort to cohort. No “explanation” of the data in Fig 4 would be very satisfying which did not succeed in attributing much of the scatter within populations to year-to-year changes in things like stream flow or ocean harvest effort, and differences between populations to things like basin size.

## 6 Discussion

For the most part, mathematical ecology has outgrown the search for “laws” analogous to those of physics. Forms such as the Beverton-Holt function for discrete population dynamics, or the logistic models for continuous population dynamics, are adopted as conventional starting points for analysis, in the same spirit that linear models or normal distributions are used to explore other kinds of data.

Almost any data analysis these days is likely to include some linear regressions, complete with *R*^2^ statistics, even when there is no particular reason to expect data to come from a truly linear relationship, or for the errors to be independent identically-distributed normal variables. From this point of view, the question of whether to start with Beverton-Holt, or Skellam, or the logistic hockey-stick, is analogous to the question of whether to plot xy-data on plain axes, or to use a log or probit transform on one or both axes.

There are, however, significant differences between the *uses* to which linear model fits and stock-production fits are typically put. The mean of a normal fit, or the slope of a linear fit, are attempts to capture some property of the “center” or “main body” of the data. The *r* and *K* parameters of a Beverton-Holt-like function, however, concern properties of the fitted curve at the fringes of the data, or in many cases well beyond them.

It is well-known that interpolation is a much safer process than extrapolation, and when a fitted model is to be used in an “interpolatory” way, for example, to make short-term forecasts, or to explore the implications of modest changes to habitat under hypothetical conditions broadly similar to historical conditions, the precise form of the fitted model is unlikely to be of great importance.

However, to estimate a true intrinsic productivity or ultimate carrying capacity from passively-observed stock-prodution data, by fitting *any* kind of stock-production model, is always an extrapolation. Statistics is not magic, and information which is not present in the data to begin with cannot be extracted from it by mathematical manipulations. In the case of stock-production:

- If the data do not include cases of low stock densities, *the data contain no intrinsic information about productivity at low densities*. Any estimate of intrinsic productivity will be driven *primarily* by *a priori* assumptions about the model form.
- If the data do not include cases of production near the carrying capacity, *the data contain no intrinsic information about carrying capacity*. Any estimate of carrying capacity will be driven *primarily* by *a priori* assumptions about the model form.
- Even if the data do include low or high stock densities, the “best” fit of a model form to the overall data need not reflect the actual behavior at these extremes very well.

This is not a problem if there are sound reasons to prefer some model form over others, or if policy conclusions are robust with respect to the choice of form. I have tried to demonstrate in this paper, however, that the space of “Beverton-Holt-like” model forms is much larger than is generally appreciated, and there are important application areas in which the choice of form matters a great deal.

## Supporting information

**S1 Appendix. Mixtures**.

**S2 Appendix. Composition Laws**.

**S3 Appendix. Stock-production functions associated with differential equations**.

